# Simulating bipedal walking using a translating center of pressure

**DOI:** 10.1101/723445

**Authors:** Karna Potwar, Dongheui Lee

**Affiliations:** Technical University of Munich (TUM); German Aerospace Center (DLR)

## Abstract

During walking, foot orientation and foot placement allow humans to stabilize their gait and to move forward. Consequently the upper body adapts to the ground reaction force (GRF) transmitted through the feet. The foot-ground contact is often modeled as a fixed pivot in bipedal models for analysis of locomotion. The fixed pivot models, however, cannot capture the effect of shift in the pivot point from heel to toe. In this study, we propose a novel bipedal model, called SLIP_*COP*_, which employs a translating center of pressure (COP) in a spring loaded inverted pendulum (SLIP) model. The translating COP has two modes: one with a constant speed of translation and the other as the weighted function of the GRF in the fore aft direction. We use the relation between walking speed and touchdown (TD) angle as well as walking speed and COP speed, from existing literature, to restrict steady state solutions within the human walking domain. We find that with these relations, SLIP_*COP*_ provides steady state solutions for very slow to very fast walking speeds unlike SLIP. SLIP_*COP*_ for normal to very fast walking speed shows good accuracy in estimating COM amplitude and swing stance ratio. SLIP_*COP*_ is able to estimate the distance traveled by the COP during stance with high precision.

## 1 Introduction

Walking is an efficient form of locomotion which allows humans to travel from one place to another. Human walking has two facets, which are reducing cost of locomotion and gait stabilization. Foot placement and orientation plays an important role in stabilizing our gait. To improve our understanding of walking and its underlying mechanism, human motion capturing and reductive modeling have been widely used [1]. Human body is a complex redundant system and reductive models or templates strip down the complex architecture of the body into simple elements and allow a computationally inexpensive way to simulate locomotion, with a certain degree of accuracy [1]. The earliest versions of such templates include the sagittal plane based inverted pendulum model (IP) [2, 3] and spring-mass model [4, 5]. The IP model assumes incompressible legs with center of mass (COM) vaulting over the legs, with a fixed foot. Due to its rigid legs, IP model provides an incorrect representation of ground reaction force (GRF) pattern, compared to that observed in humans. To overcome this drawback of the IP model, some templates include a springy leg to obtain more accurate estimates of walking gait trajectories [6, 7, 8, 9, 10]. One such model is the sagittal plane based SLIP model [10]. Due to SLIP’s compliant legs it is able to generate the COM trajectory and GRF pattern during walking to that observed in human walking. The steady state solutions of SLIP also suggests that walking is one of the many domains of different locomotion patterns generated. Hence, it is important to narrow down the parameters in a bipedal model pertaining only to the domain of walking.

IP model and SLIP model assume a fixed pivot during stance as discussed above, which is essentially the mean position traveled by COP during stance. In a study to analyze treadmill walking^1^, SLIP fairly estimates COM trajectory at 1m/s, while failing to estimate at other walking speeds [11]. They also mention that due to the fixed pivot of the SLIP, the model has to be simulated at a steeper TD angle compared to human walking. This characteristic of SLIP might be restricting its predictive capabilities at slower and faster walking speed. During walking, COP of a particular foot travels approximately a distance of a foot length and the COP progression velocity depends on speed of walking [12, 13, 14]. To accommodate the mechanical consequences of COP progression, a few bipedal models were developed for running [15, 16] and walking [17, 18]. Bullimore et al. [16] take into consideration the change in TD and LO angles caused by COP translation, called POFT (point of force translation), in the conventional spring mass model for running. They use a constant velocity based COP progression model without considering the acceleration of the COP during stance. They observe similar COM trajectories and GRF patterns as in human running but the model shows drastic decrease in spring stiffness. Lee et al. [17] use a translating point of force application (PFA) in an IP model for walking and show that the error in vertical displacement of the COM predicted by the model increases from 111% to 240%, as walking speed increases from 0.5 m/s to 2.5 m/s respectively. However, these errors were considerably lesser compared to IP model with a fixed pivot. Miff et al. [18] show that during walking vertical excursion of the trunk is dependent on the foot rocker radius in the rocker based IP model. IP models in the above studies consider only the single stance vaulting of the COM but not the foot impact and double stance phase of walking. We need a bipedal model which can simulate COP progression along with single/double stance, COM trajectory and GRF patterns.

The objective of this study is to check, if addition of a COP progression model would improve SLIP model’s performance at slower and faster walking speeds. Instead of using a predefined leg stiffness [11], we optimize our spring stiffness. We use the relation between TD angle, walking speed and COP speed obtained from existing literature so as to make model-experiment comparison. We develop a generic model called SLIP_*COP*_ (Fig. 1) with COP progression considering two modes of COP translation during stance: one with a constant COP speed and the other accelerated COP. We include the constant velocity COP progression model so as to compare our model with previous similar models. We make inter-model comparisons between SLIP and SLIP_*COP*_ for various walking parameters to analyze the results qualitatively and quantitatively. Subsequently, we compare the two models with real walking scenarios to assess the quality of our solutions.

**Figure 1:**
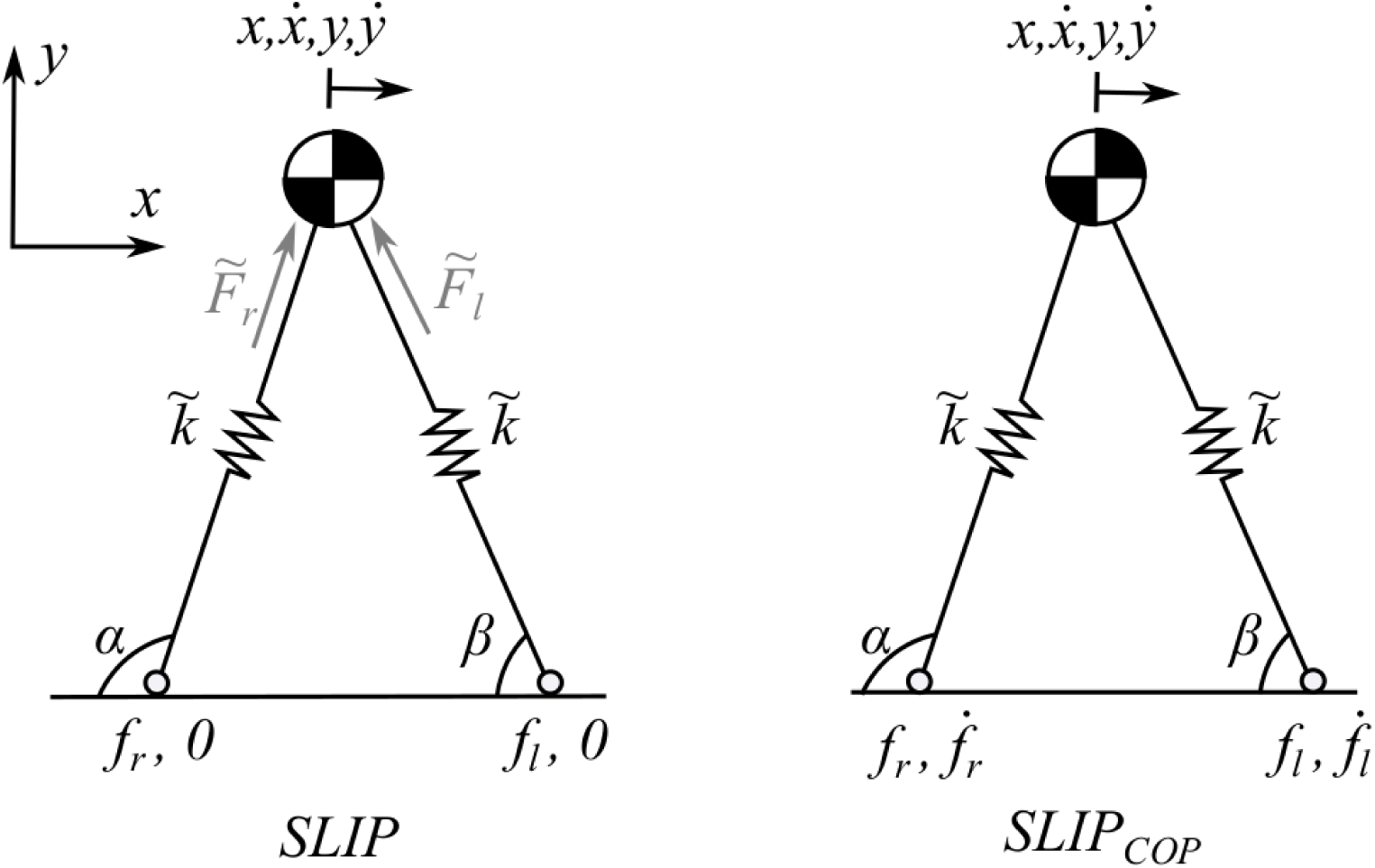
Diagram showing the two models, (Left) SLIP and (Right) SLIP with translating COP (SLIP_*COP*_)with COM and COP coordinates in the sagittal plane. Subscripts r and l stand for right and left leg respectively.

## 2 Method

We simulate the two models, SLIP and SLIP_*COP*_, as seen in Fig. 1. The position and velocity of right and left COP are denoted as 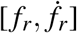 and 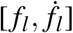 respectively. For SLIP, 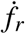 and 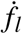 are always 0 due to a fixed pivot. Like the conventional SLIP, SLIP_*COP*_ consists of a COM attached with two springy legs. The legs are considered massless and a swinging leg can be ignored. As illustrated in Fig. 2, COM state at apex is described by 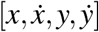 and at TD the leg makes an angle of *θ*_*o*_. Both models are simulated in the sagittal plane. We non-dimensionalize the equations of motion to develop generic models catering to humans with different anthropometric measurements [19, 20, 21]. Force experienced by the COM before non-dimensionalization in the forward and vertical direction is given as

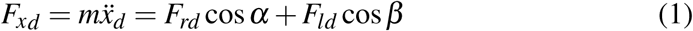

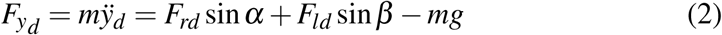

where *F*_*rd*_ = *k*(*L*_*o*_ *−L*_*rd*_) is the GRF in the right leg, *m* is the mass, *k* is the spring stiffness, *g* acceleration due to gravity, 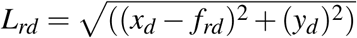 is the length of the right leg in stance, *L*_*o*_ is the uncompressed leg length, the subscript *l* and *r* refer left and right leg, and the subscript *d* means dimensionalized. Upon non dimensionalizing eqns.(1)(2), the time-dependent terms are divided by 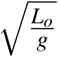, distance terms by uncompressed leg length *L*_*o*_ and divide the equations throughout by *mg* [20, 10]. After non-dimensionalization the force experienced by the COM is given as

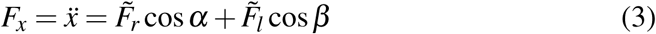

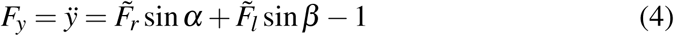

Where 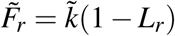 (see Fig. 1),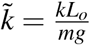 is the relative stiffness of the legs. At TD, the leg angle reorients to *θ*_*o*_ and at lift off (LO) occurs when the GRF becomes 0.

**Figure 2:**
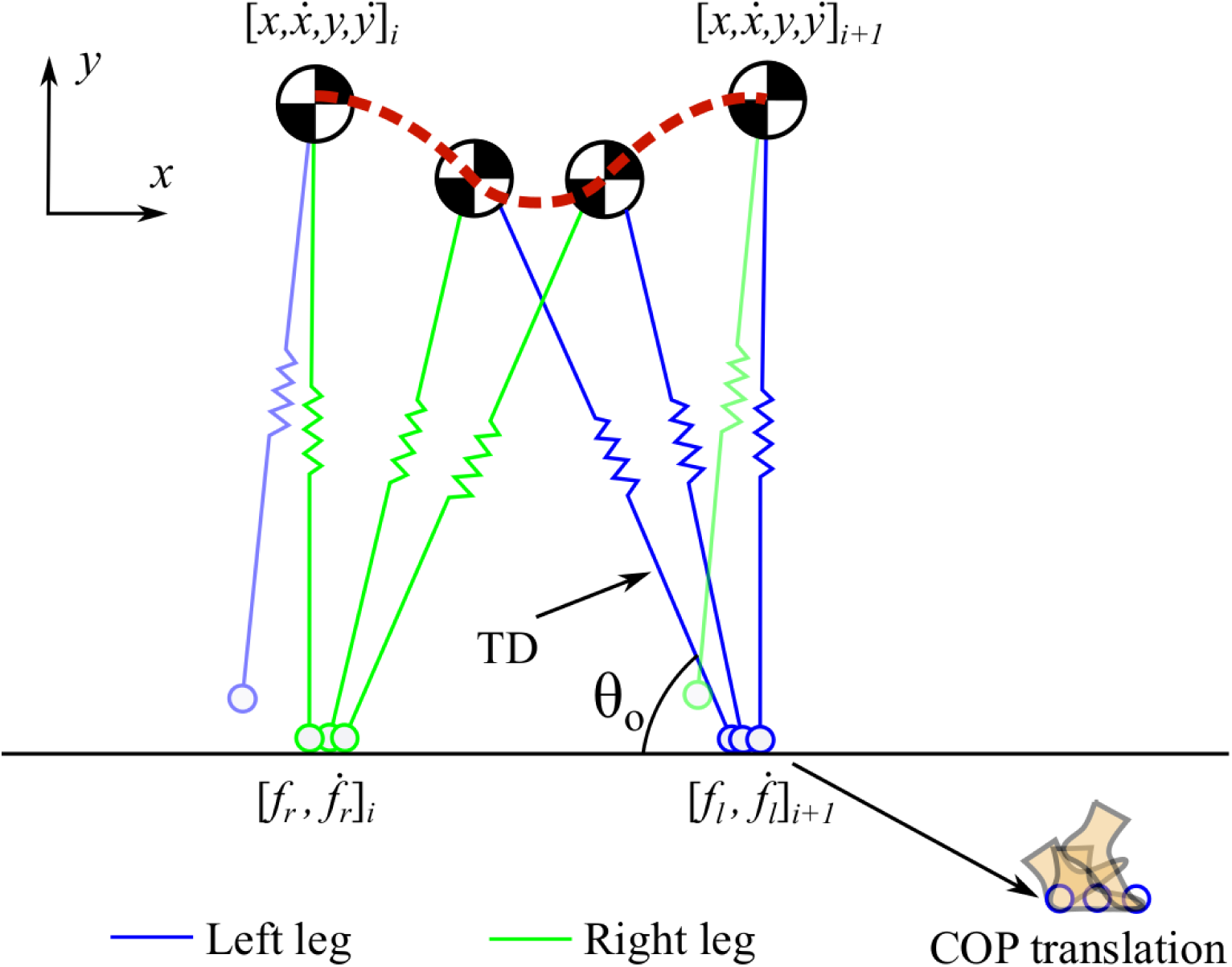
A limit cycle of the translating COP model. The model starts at the apex *i* and attains the consecutive apex *i* + 1, while the COP translates along the ground.

### 2.1 Gait parameter relations

In order to restrict our model’s solution search within the walking domain, we use the relation between walking speed, TD angle and COP speed obtained through existing literature [11, 12] (see Table 2). The lower and upper bound for the COP progression velocity are the minimum and maximum speed of the COP during experimental walking. The COP model during stance is described as the function of the GRF in the fore-aft direction as

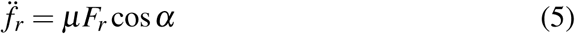

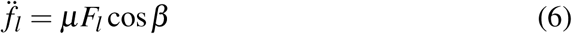

**Table 1:**
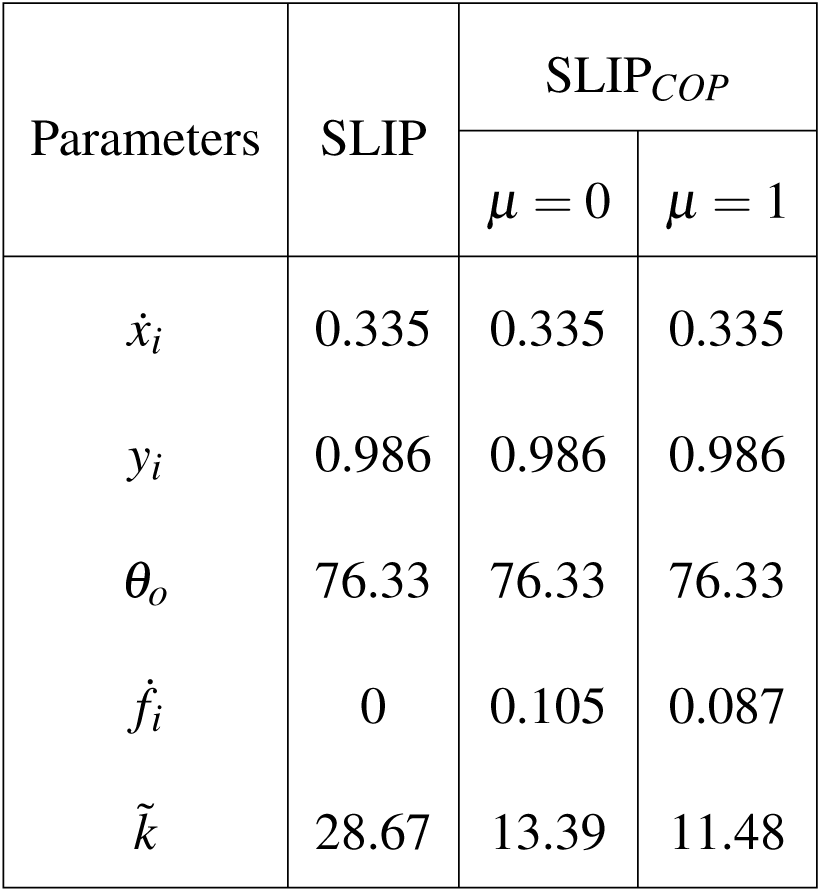
Simulation parameters for results in Figure 5. 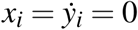.

| Parameters | SLIP | SLIP<sub>COP</sub> |  |
| --- | --- | --- | --- |
| | | $\mu = 0$ | $\mu = 1$ |
| $\dot{x}_i$ | 0.335 | 0.335 | 0.335 |
| $y_i$ | 0.986 | 0.986 | 0.986 |
| $\theta_o$ | 76.33 | 76.33 | 76.33 |
| $\dot{f}_i$ | 0 | 0.105 | 0.087 |
| $\tilde{k}$ | 28.67 | 13.39 | 11.48 |

**Table 2:**
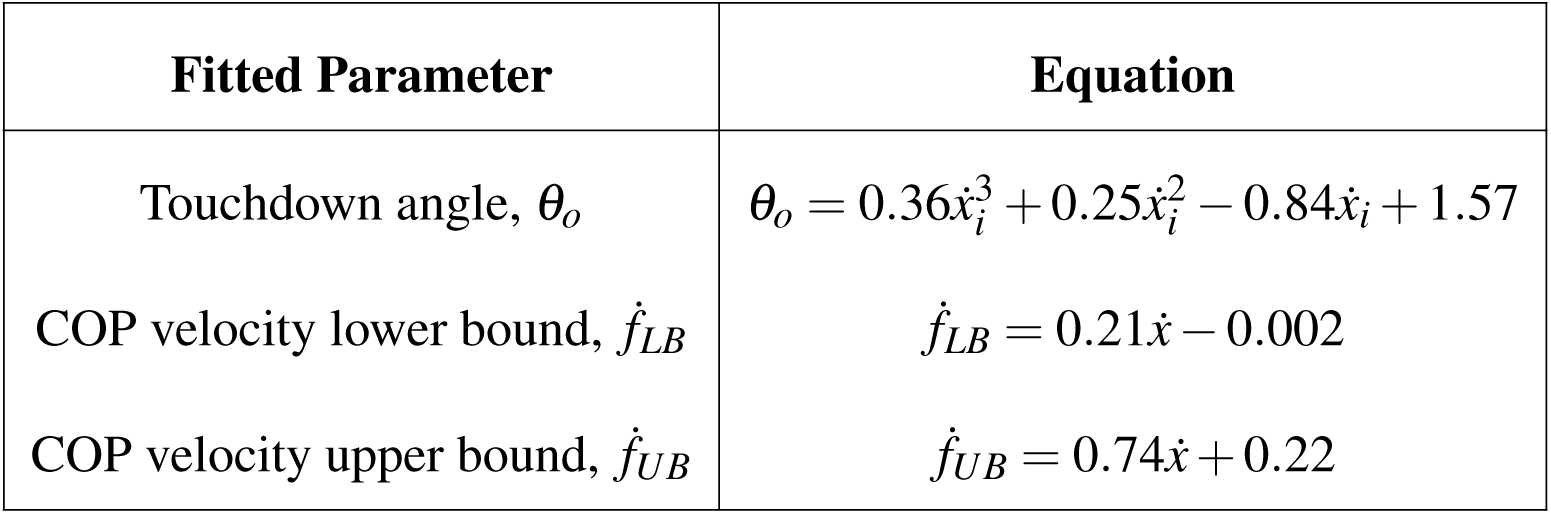
Parameter relations obtained from physiological data[11, 12]. Subscript *LB* and *UB* stand for lower bound and upper bound respectively.

Especially, two modes of this translating COP model are considered: one considering effect of a constant COP speed during stance (*μ* = 0) and the other as weighted function of the GRF during stance (*μ* = 1).

We obtain steady state solutions of the two models by optimizing their parameters to generate a limit cycle. To generate a limit cycle we consider the apex to apex state errors. The state of the model at apex is completely described by its relative horizontal distance between COM and COP denoted by (*x − f*), horizontal velocity 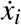 apex height *y*_*i*_, vertical velocity 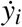 and COP velocity 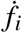 To obtain a limit cycle, we calculate the stride to stride error for consecutive apex states (*i* and *i* + 1) using a 5-dimensional nonlinear Poincaré return map [22]. The initial apex state and final apex state of the model are given as 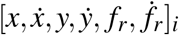 and 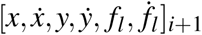 respectively. For a given set of 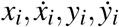, we optimize relative stiffness 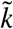, right COP speed 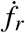 and left COP speed 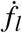 to get a limit cycle as shown in the Algorithm 1. At the start of simulation, the foot is placed at the origin with the COM directly above it.

#### Algorithm 1

Algorithm to obtain a limit cycle

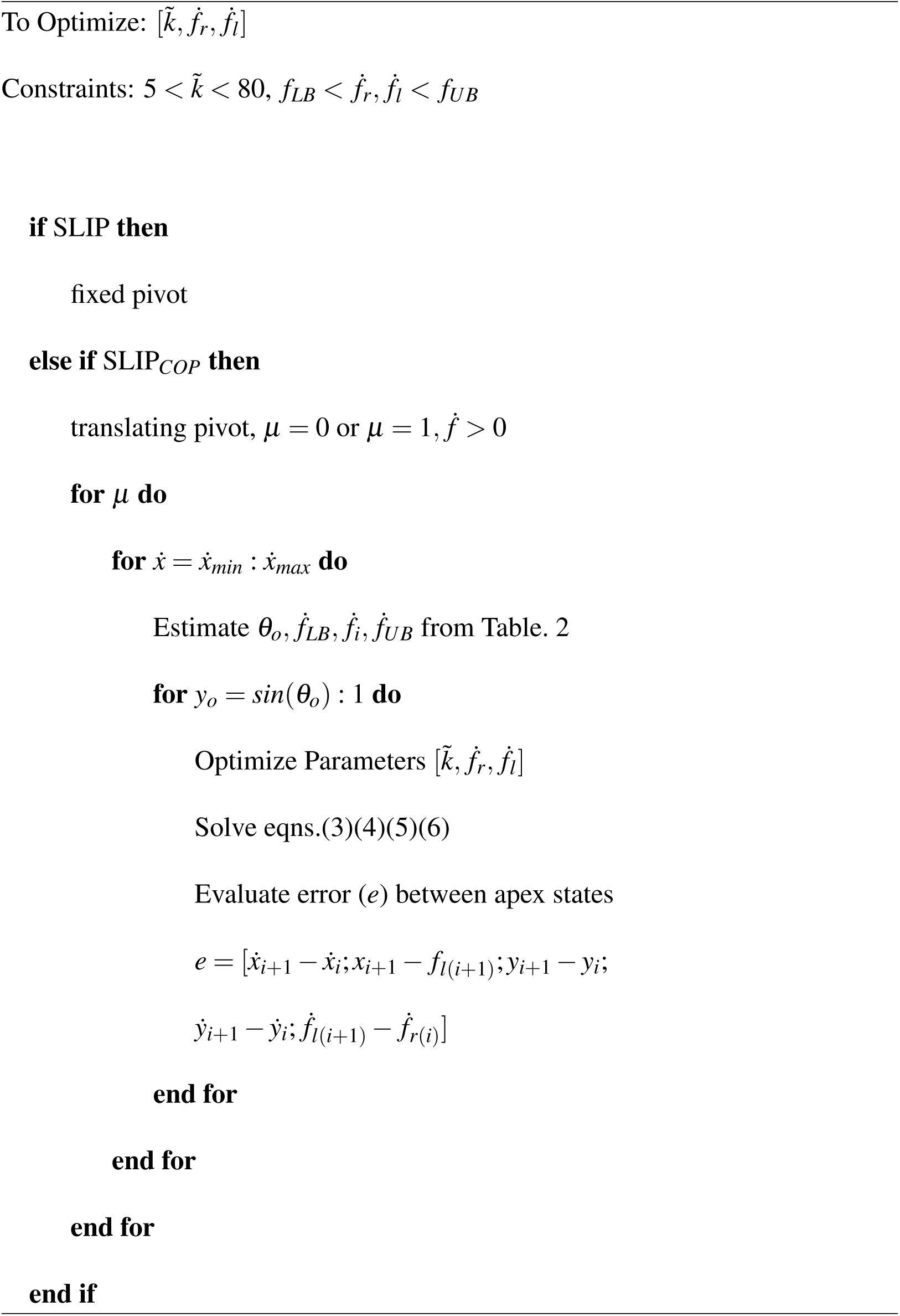

## 3 Results

Firstly, we compare COM, COP and GRF trajectories for the two models (SLIP, SLIP_*COP*_) for a given set of optimized parameters (see Table 1) to assess the qualitative nature of the solutions. Fig. 4a & b are GRF and COP trajectories for individual legs. GRF pattern in vertical and horizontal direction resemble that of experimental walking[10, 11, 23]. SLIP_*COP*_ shows a lower vertical GRF value at mid-stance (F_*y*_), for both of the COP modalities (*μ* = 0 and 1) compared to SLIP. At *μ* = 0 the COP translates with constant speed and at *μ* = 1 the speed results in a U-shape profile which correlates to the horizontal GRF (F_*x*_) (Fig. 4b). The shape of the COP speed trajectory for *μ* = 1 resembles the COP speed trajectory of human walking. As seen in Fig. 4c, SLIP_*COP*_ has higher COM amplitude and horizontal distance travelled than SLIP. This result, as expected, is a consequence of COP progression. A higher gait distance for *μ* = 1 is due to a larger average COP speed (see Fig. 4b) compared to at *μ* = 0.

**Figure 3:**
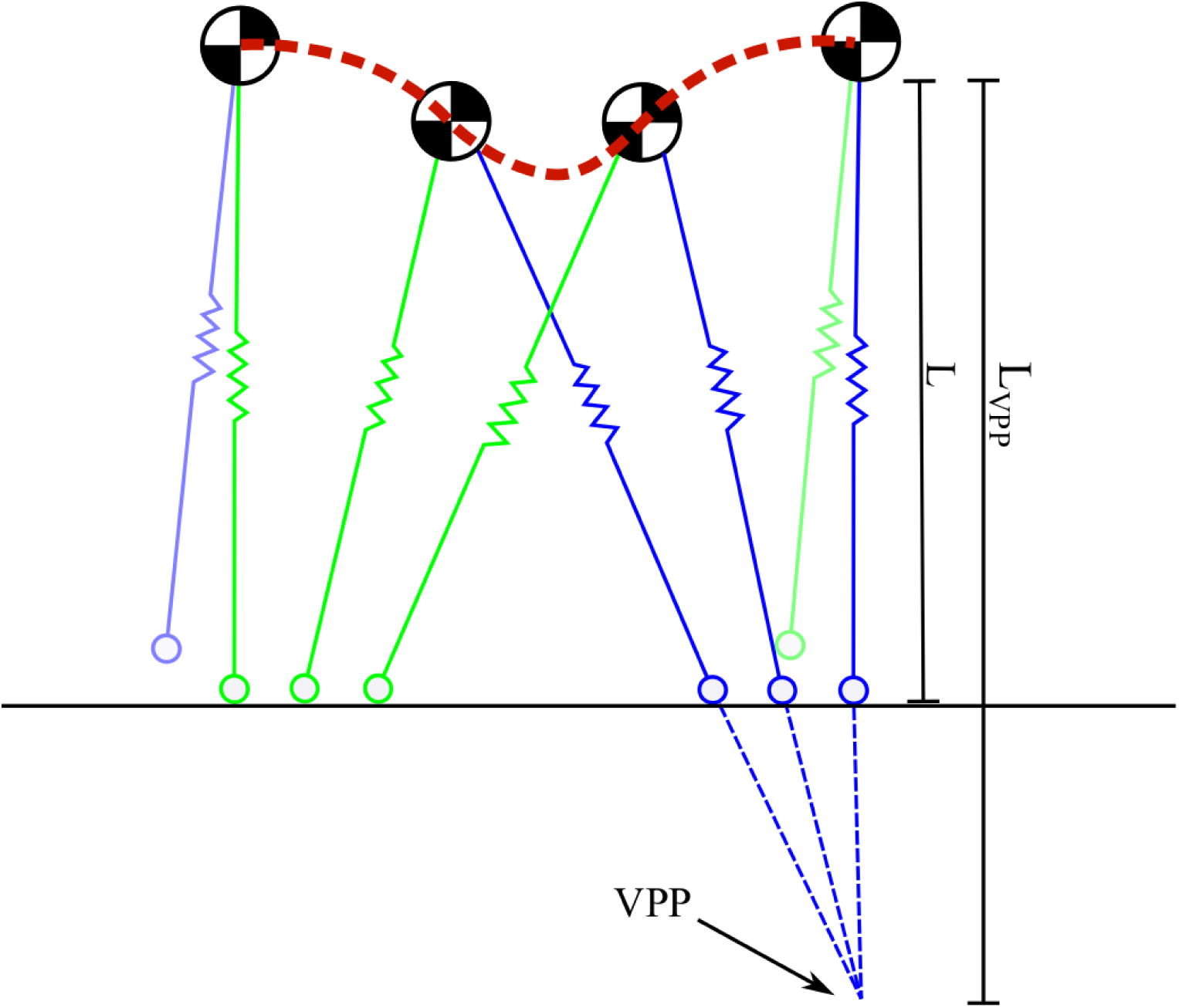
Translation of the COP in SLIP_*COP*_ leads to a virtual pivot point (VPP) under the surface. *l*_*v*_ is the extended length where *l*_*v*_=1.8*l* [18] during walking.

**Figure 4:**
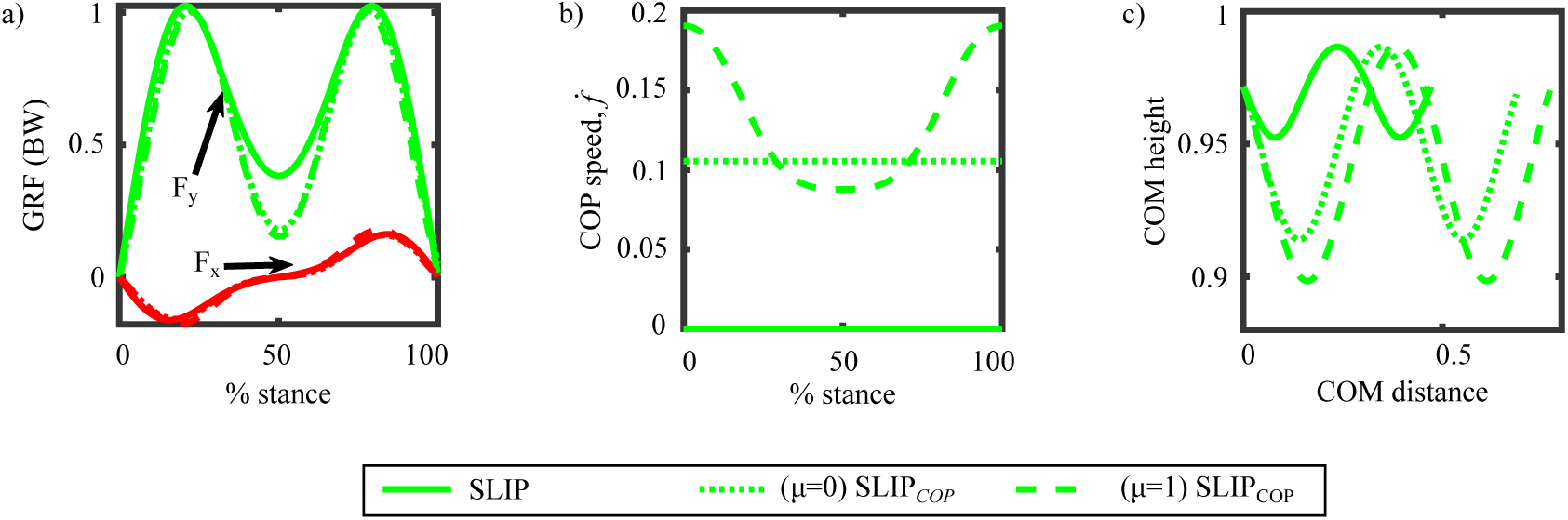
GRF, COP speed and COM trajectory plotted for SLIP and SLIP_*COP*_ (*μ* = 0 and *μ* = 1) with parameters in Table 1.

We make inter-model comparisons at a non-dimensionalized speed 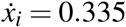 and *θ*_*o*_ = 76.33*°*. We compare the steady state solutions, at the mentioned speed and TD angle, obtained by varying *y*_*i*_ *∈* (sin(*θ*_*o*_), 1). We discard solutions at *y*_*i*_ = *sin*(*θ*_*o*_) and *y*_*i*_ = 1. Because at *y*_*i*_ = sin *θ*_*o*_ the apex height will be equal to the COM height at TD, which is physically impossible. And at *y*_*i*_ = 1, the system will be under free fall as the leg will be at its natural uncompressed length suggesting no foot contact with the ground. As seen in Fig. 5a for SLIP, increase in *y*_*i*_ leads to increase in 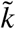 from 22.75 to 29.54. SLIP_*COP*_ for both COP modalities shows a considerably lower and constant stiffness for all values of *y*_*i*_. We expected lower stiffness for SLIP_*COP*_ because of the leg lengthening that occurs due to a virtual pivot point generated as shown in Fig. 3. During walking, stride length is approximately twice the value of step length as seen in Fig. 5c & e. A higher value for step lengths for SLIP_*COP*_ is observed: e.g. a value of 0.46 at *y*_*i*_ = 0.99 with *μ* = 1. For walking, cadence *c* and step length *s*_*l*_ are related to walking speed as *v* = (*c*)(*s*_*l*_) [18]. We see the effect of this hyperbolic relation between cadence *c* and step length *s*_*l*_ in the plots Fig. 5c & d. The swing/stance duration ratio is around 0.4 for walking, and SLIP achieves this ratio as *y*_*i*_ approaches 1 (see Fig. 5f). SLIP_*COP*_ shows a reduced swing/stance duration time which occurs due to its increased stance time. This increase in stance time occurs as a result of the COP progression which we expected [16].

**Figure 5:**
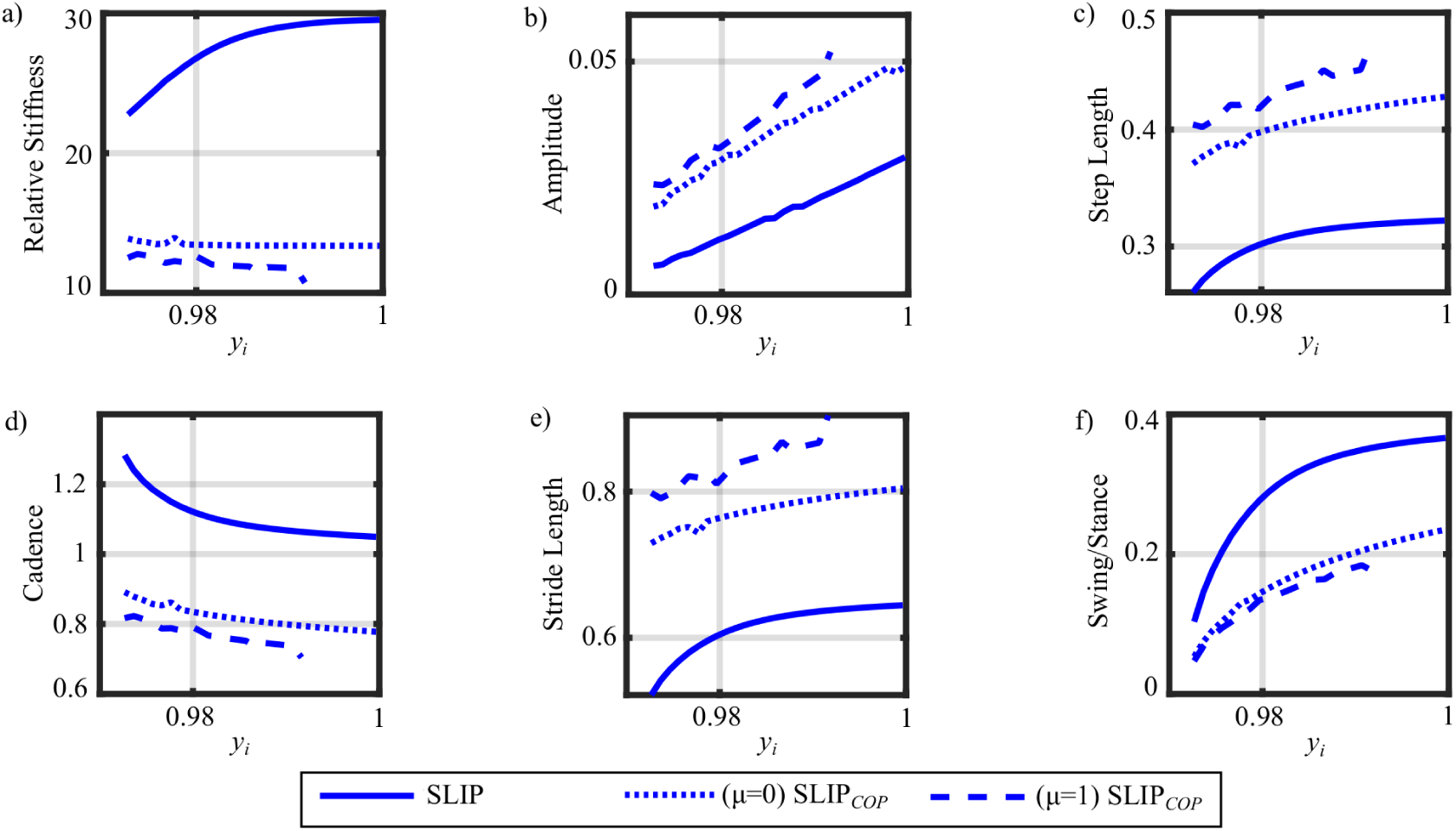
Plotting temporal and distance variables for different values of *y*_*i*_ as at an apex speed 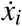 of 0.335 and *θ*_*o*_ of 76.33*°*.

The reliability of the two models is tested by making model-experiment comparisons. In particular, we compared the mean error in between model and experiment data for the following parameters: vertical COM amplitude *a*, swing/stance ratio, walking speed *v*, virtual pivot point (VPP) length factor *γ*, COP speed and distance (*D*_*COP*_) travelled as shown in Fig. 6. We use the following equations from existing literature for experimental walking.

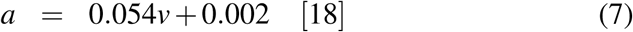

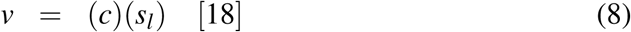

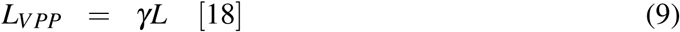

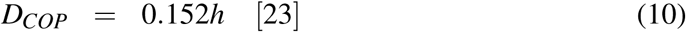

where *c* is cadence, *s*_*l*_ is step length, *L*_*V PP*_ is distance between COM and virtual pivot point, *γ* is the VPP factor usually around 1.8, *D*_*COP*_ is distance travelled by the COP during stance and *h* is the height of the human. Winter et al. [23] provided measures of ratios of different body parts with respect to human height. Such a ratio for foot measure is shown in eq.(10). Swing/stance ratio is calculated by dividing the swing time of a particular leg by its stance time during a gait cycle. To compare the models with experiment data of adults we dimensionalize the parameters. Walking speed is dimensionalized by multiplying 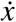 with 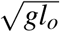, where *l*_*o*_=1m is the uncompressed leg length. In Fig. 5, we compared solutions at every apex *y*_*i*_ at 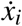 f 0.335 and *θ*_*o*_ = 76.33*°*. In Fig. 6 we calculate the average of all limit cycle solutions for all dimensionalized apex speeds to make model-experiment comparison.

**Figure 6:**
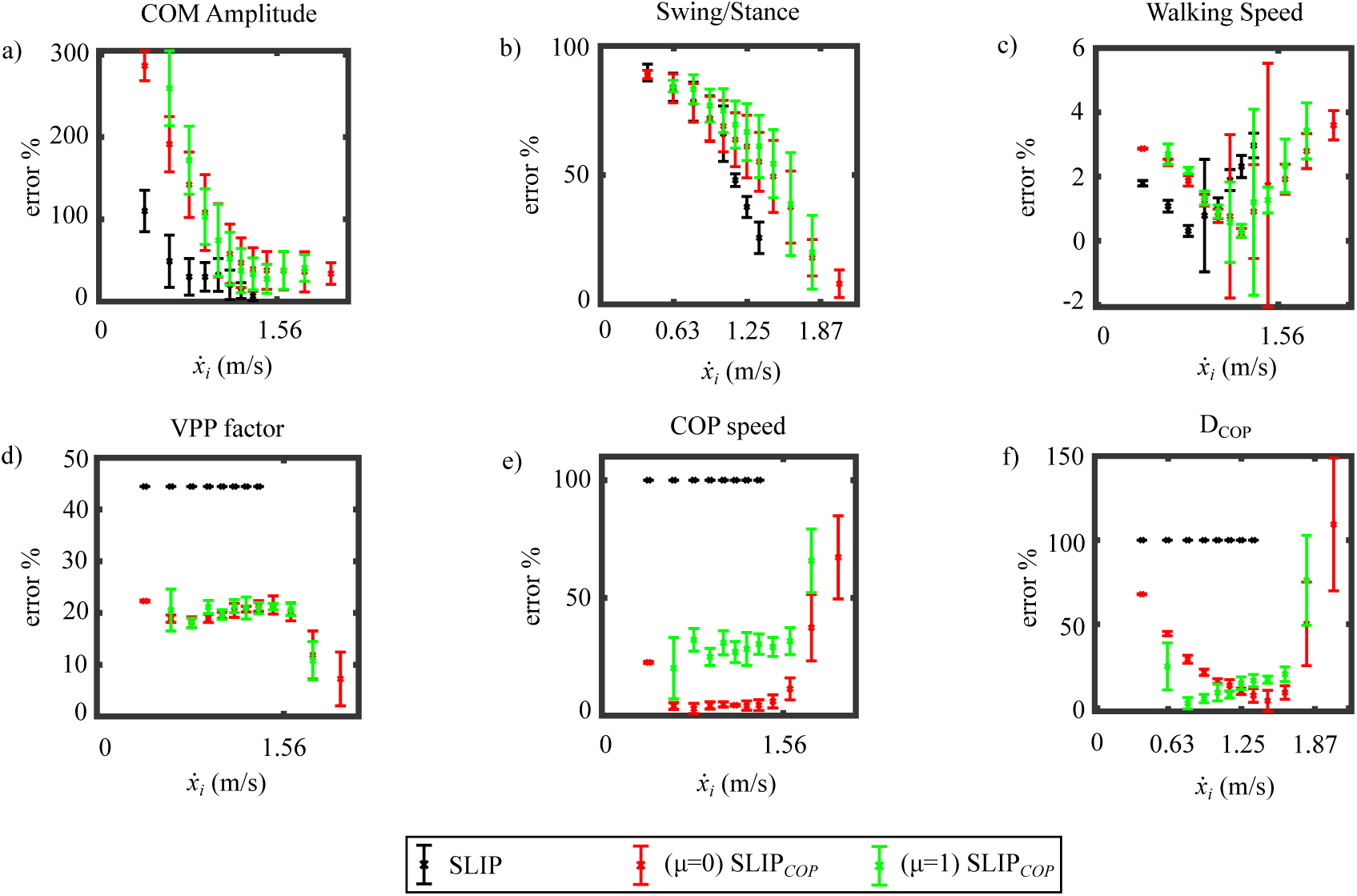
Mean errors of the obtained model parameters (amplitude, swing/stance duration ratio, average walking speed, VPP factor, average COP speed and COP distance) with respect to experimental human data.

One of the objectives of our study was to to see if the relation among TD angle, walking speed and COP speed provides steady state solutions for the given adult speed range^2^. We get steady state solutions for very slow to slow walking speeds for SLIP and very slow to very fast walking speeds for SLIP_*COP*_. For SLIP_*COP*_ with both *μ* values, we see a larger error for COM amplitude and swing/stance ratio at very slow to normal walking speeds, compared to SLIP. But the error decreases as the walking speed increases. SLIP estimates COM amplitude much better than SLIP_*COP*_ at lower speed ranges because its optimized spring stiffness values lie close to human leg stiffness. Fig. 6b illustrates the error of swing/stance ratio. At lower walking speed, both models perform similarly. As walking speed increases, the errors for both models decrease. At the normal walking speeds SLIP provides less error than SLIP_*COP*_; at 1.36 m/s, 25.72% error for SLIP, 55.17% for SLIP_*COP*_ (*μ* = 0), and 61.13% for SLIP_*COP*_ (*μ* = 1). For higher walking speeds, SLIP_*COP*_ outperforms (error below 25%) SLIP, which fails to find a solution.

The concept of virtual pivot point (VPP) is illustrated in Fig. 3 and expressed in eq.(9). In Fig. 6d the VPP factor (*γ*) is compared with the physiologically measured value of 1.8 provided by [18]. As expected, SLIP model provides *γ* which is equal to 1 under all scenarios because of its fixed pivot. SLIP_*COP*_ (*μ* = 0, 1) provides more accurate estimates of *γ* with approximately 20% error. VPP is a good metric to measure the effectiveness of our COP progression model. The error remains constant at 20% for most of the speed range but decreases to 10% at very fast speed ran (see Fig. 6e). *D*_*COP*_ is calculated, for an adult with an average height of 1.7 meters [11] with an uncompressed leg length, *L*_*o*_ = 1*m*. In Fig. 6f we observe, SLIP_*COP*_ shows quite a low error except at very slow and very fast walking speeds: the least error of 4% at 0.75 m/s with *μ* = 1 and 7% at 1.4m/s with *μ* = 0.

## 4 Discussion

Through this study we compared the effect of adding a translating COP to the conventional SLIP model. The motivation behind using a translating COP was to simulate the effects of the heel-to-toe pivoting of the foot during stance phase in human walking. One of the objectives of our study was to improve the predictive capabilities of SLIP for a wider speed range^2^ when comparing with experimental data. Utilizing experimental data (of the relation among walking speed, TD angle, and COP speed) enables us to obtain limit cycle solutions at these speed ranges. The relation between TD angle and walking speed also suggests that as the walking speed approaches 0, TD angle approaches 90 degrees (erect standing), which can be considered as decent validation of our walking speed and TD angle relation. This shows that as the walking speed approaches a lower value step length approaches 0. Lipfert et al. [11] showed that to obtain similar walking dynamics at a given walking speed, the SLIP model was simulated at a steeper angle because of premature lift off at the correct TD angle. With the relation between TD angle and walking this limitation is overcome. On the other hand, provision of COP speed-walking speed relation leads to increase in stance time. When comparing the 2 models at similar optimized state variables, we see an increase in gait distance for SLIP_*COP*_ because of increase in its stance time. To comment upon gait distance estimation of the 2 models we dimensionalize the result in Fig. 4c with a leg length *L*_*o*_ = 1*m* and speed of 1m/s so as to compare to a previous model-experiment study using SLIP [11]. Upon comparing, the COM trajectories in Fig. 4c, we observe that SLIP_*COP*_ estimates the gait distance with an error of m and SLIP with an error of 0.25 m [11]. One of the reasons for this underestimation by SLIP could be its fixed pivot point. Although we used constant stiffness in our models, we preferred optimizing the spring stiffness rather than using predefined leg stiffness [11]. This was done because the relation among TD angle, walking speed, and COP speed could affect the optimal value of stiffness. We understand that the human walking gait is a consequence of the stabilization occurring at the foot. This has led us to put more emphasis on the relation among TD angle, walking speed, and COP speed rather than on leg stiffness as done by Lipfert and colleagues. Although human muscular strength determines the flexion and extension of our lower limbs during walking, it is difficult to measure this strength just by observation. Through inverse dynamics we can utilize observable kinematic and dynamic characteristics to understand more about the functioning of human walking. The COP speed trajectory for the accelerated COP modality shows a similar trend in COP speed as shown in the study by Cornwall and colleagues [25], with high speed at initial contact phase, lower speed at mid stance and higher speed at TD. This U-shape speed profile correlates with the horizontal COM acceleration F_*x*_ because COM decelerates in the first half of stance phase and then accelerates in the next half. This suggests that our weighted function approximates the acceleration of the COP quite well.

In Fig. 4a & c we see the effects of a translating COP which leads to higher variation in vertical GRF and COM amplitude respectively, compared to SLIP. Such behavior was also observed in the IP model [17] and POFT model [16], where the addition of a translating COP increases the vertical displacement significantly. Bullimore et al. [16] also mention that due to a translating COP the stance time for a leg increases which we also observe in SLIP_*COP*_. They also mentioned that addition of a COP progression model decreases the spring stiffness of the model. The decrease in spring stiffness can be explained by eqns.(11)(12)(13). In Fig. 4a, F_*y*_ at mid stance (apex) is lower for SLIP_*COP*_ than SLIP. At apex the COM experiences centripetal acceleration due to its weight and spring force. Hence, upon referring Fig. 2, 3 and Table 2, the force balance for SLIP and SLIP_*COP*_ at apex is

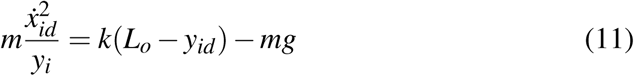

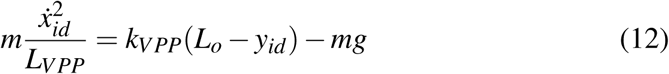

Subtracting eqns.(12) from (11) we get,

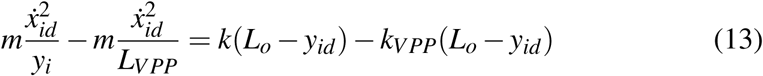

As both sides of the eqn.(13) are positive with *y*_*i*_ *< L*_*V PP*_, this implies *k > k*_*V PP*_. To reduce the vertical displacement occurring due to reduced spring stiffness, Bullimore et al. [16] added a constraint on the vertical movement of the COM. Constraining the vertical displacement for our models resulted a difficulty to find limit cycle solutions and hence we relaxed this constraint. Lee et al. [17] showed that with increasing walking speeds the error in COM vertical displacement increases when compared to experimental data. We observe a decrease in error for COM vertical displacement, at *μ* = 0 and 1, with increasing walking speeds which shows the effectiveness of our bipedal model. The IP model in the above studies was simulated only for single stance which could have limited its predictive nature unlike the SLIP and SLIP_*COP*_.

One of the characteristics of walking is the relation between cadence and step length represented by eqn.(8). We obtain quite low errors for both models for walking speed using eqn.(8) as seen in Fig. 6c. We observe an increase in step length and decrease in cadence for SLIP_*COP*_ which was expected in our study. With COP progression the distance between consecutive heel strikes increases subsequently increasing the step length. This in turn reduces cadence (refer eqn.(8)). As discussed before, due to COP progression we have an increase in stance time which is also responsible for decrease in cadence because is defined as steps per min. One more factor that is characteristic of a progressive COP model is the generation of a virtual pivot point as discussed above (Fig. 3). To the best of our knowledge, there exists no study with SLIP model that has estimated the VPP factor *γ*. The average *γ* value with our proposed SLIP_*COP*_ is around 1.4 with a 20% mean error, where the range of *γ* is between 1.33 and 2.1. To put the value of *γ* into perspective, we evaluate the distance travelled by the COP during stance. As COP travels approximately a foot length [12], we evaluated the COP distance D_*COP*_ for our speed range. The accelerated COP modality shows a considerably lower error values than the constant velocity modality for very slow to normal walking speeds. One of our objectives was to differentiate between the two COP modalities *μ* = 0 and 1. With D_*COP*_ we see that for very slow to slow speeds *μ* = 1 provides better estimation and for normal walking speeds *μ* = 0 is better. Overall the accelerated model shows lower error value for D_*COP*_ for majority of speeds suggesting its reliability over the constant velocity modality.

We propose a bipedal spring mass model utilizing the COP translation observed during human walking. We compare this model with the SLIP model with respect to human walking data. We observe that the SLIP and SLIP_*COP*_ show pretty high error estimates for COM vertical amplitude and swing/stance ratio at very slow to slow walking speeds. At normal to very fast walking speeds, we see the benefits of the SLIP_*COP*_ as it not only provides limit cycle solutions for these speed zones but also considerably decreases error in predicting COM amplitude and swing/stance duration ratio. SLIP_*COP*_ is able to reproduce a symmetrical COP speed profile in the fore-aft direction. The distance traveled by the COP for the two COP progression modes at normal walking speeds concurs with distance traveled by the COP during human walking and can be considered as a substitute for an ankle based walking model. This pilot study on using a translating COP based SLIP model takes into consideration the fact that COP movement is closely related to the GRF force in the horizontal direction.

In the future work, this model can be further developed by utilizing actual COP data from human walking, which could enhance our models capabilities from the point of view of simulating slow to normal walking speeds. This study will be also undertaken from the point of view of assessing gaits in people with movement disorder such as Cerebral Palsy, Stroke and Parkinson’s. People with such movement disorders often portray unequal strength in their legs. This affects their walking style, foot placement consequently affecting their COP dynamics. Developing our COP model towards adapting it to assess these movement disorders would help us understand the difference between healthy and impaired walking styles.

## Supporting information

Cover Letter

## List of abbreviations

## Abbreviations

GRF: Ground reaction force
SLIP: Spring loaded inverted pendulum
COM: Center of mass
COP: Center of pressure
TD: Touch down
IP: Inverted pendulum
POFT: Point of force translation
LO: Lift off
VPP: Virtual pivot point

## Acknowledgements

We would like to thank Dr. Leif Johannsen and Dr. Ing. Matteo Saveriano for their inputs to our research.

## Author contributions

Karna Potwar conducted the research, dealing with generating the hypothesis as well as simulating and analyzing the bipedal walking model. He also documented and edited the manuscript. Dongheui Lee supervised this research and critiqued the manuscript.

## Footnotes

1 Subjects walked at speeds of (0.52,1.04,1.55,2.07 and 2.59) m/s in this experiment

2 We classify walking speeds as very slow (0.7-1.12 m/s), slow (1.12-1.31 m/s), normal (1.31-1.58 m/s), fast (1.58-1.76 m/s) and very fast (1.76-2.19 m/s) based on the classification provided by [24].

